# Genome analysis of the esca-associated Basidiomycetes *Fomitiporia mediterranea*, *Fomitiporia polymorpha*, *Inonotus vitis*, and *Tropicoporus texanus* reveals virulence factor repertoires characteristic of white-rot fungi

**DOI:** 10.1101/2024.06.09.598094

**Authors:** Jadran F. Garcia, Rosa Figueroa-Balderas, Gwenaëlle Comont, Chloé E. L. Delmas, Kendra Baumgartner, Dario Cantu

## Abstract

Some Basidiomycete fungi are important plant pathogens, and certain species have been associated with the grapevine trunk disease esca. We present the genomes of four species associated with esca: *Fomitiporia mediterranea*, *Fomitiporia polymorpha*, *Tropicoporus texanus*, and *Inonotus vitis*. We generated high-qualityphased genome assemblies using long-read sequencing. The genomic and functional comparisons identified potential virulence factors, suggesting their roles in disease development. Similar to other white-rot fungi known for their ability to degrade lignocellulosic substrates, these four genomes encoded a variety of lignin peroxidases and carbohydrate-active enzymes (CAZymes) such as CBM1, AA9, and AA2. The analysis of gene family expansion and contraction revealed dynamic evolutionary patterns, particularly in genes related to secondary metabolite production, plant cell wall decomposition, and xenobiotic degradation. The availability of these genomes will serve as a reference for further studies of diversity and evolution of virulence factors and their roles in esca symptoms and host resistance.

## Introduction

The majority of wood-colonizing species that cause grapevine trunk diseases belong to the Ascomycota phylum, with only a few Basidiomycetes species confirmed as pathogens primarily of esca (Del Frari et al. 2021; Fischer and González García 2015; Brown et al. 2020; Fischer 2000, 2002; Martín et al. 2022). Whereas the primary pathogens of esca were long thought to be Ascomycetes *Phaeomoniella chlamydospora* and *Phaeoacremonium minimum*, various members of the Hymenochaetaceae family, including species like *Fomitiporia mediterranea* and *Inonotus* spp., have been isolated in esca-symptomatic vines in vineyards around the world (Fischer and González García 2015; Brown et al. 2020; Fischer 2002; Guerin-Dubrana et al. 2019; Pacetti et al. 2022; Mundy et al. 2020; Cloete et al. 2015b, 2015a; Moretti et al. 2021). Brown et al. (Brown et al. 2020) isolated multiple Basidiomycete species from vineyards with esca in California and Texas, and described novel species *Fomitiporia ignea*, *Inonotus vitis*, and *Tropicoporus texanus*. *Tropicoporus texanus* was capable of causing symptoms independently, without co-inoculation with *P. chlamydospora*. Moreover, leaf symptoms resembling those of esca were more prevalent when *F. polymorpha* was co-inoculated with *P. chlamydospora* than when either were inoculated alone (Brown et al. 2020).

The ability of these fungi to colonize the wood and influence disease progression is determined by their functional arsenal. Wood-rotting Basidiomycetes are commonly categorized as either white- or brown-rot fungi, based in part on the type of residue they leave after degrading their host’s wood (Worrall et al. 1997; Sista Kameshwar and Qin 2018; Eichlerová and Baldrian 2020). White-rot species can degrade all of the main components of the wood (cellulose, hemicellulose, and lignin), whereas brown-rot species degrade cellulose and hemicellulose, leaving a modified lignin residue of brown color (Moretti et al. 2021; Worrall et al. 1997; Sista Kameshwar and Qin 2018; Eichlerová and Baldrian 2020; Zhou et al. 2014). These differences are attributed to the plant cell wall degrading enzymes (CWDEs) they produce, to the activities of the encoded enzymes *in planta*, to their non-enzymatic wood-decay capabilities, and also to the condition of the woody substrate they colonize. White-rot fungi are characterized by encoding lignin-oxidizing and lignin-degrading enzymes, which are grouped into the Auxiliary Activity (AA) family of the Carbohydrate-Active Enzymes (CAZymes) (Sista Kameshwar and Qin 2018). Additionally, white-rot fungi secrete a more diverse array of Glycoside Hydrolases (GH) than brown-rot fungi. To possibly compensate for the lower diversity of GH, brown-rot fungi secrete their GHs in greater abundance than their white-rot counterparts (Presley et al. 2018).

In addition to CAZymes for the breakdown of polysaccharides in the plant cell wall (Sista Kameshwar and Qin 2018; Kameshwar and Qin 2016), Basidiomycetes utilize other enzyme groups. Among them, peroxidases are used for decomposing structural components like lignin (Kameshwar and Qin 2016; Manavalan et al. 2015; Ayuso-Fernández et al. 2019). Cytochrome P450 enzymes support multiple primary and secondary metabolism, including the degradation of xenobiotic compounds (Morel-Rouhier 2021; Syed et al. 2014) and competition with other organisms (Riley et al. 2014; Castaño et al. 2022). Lastly, transporters facilitate the intake and secretion of various molecules and enzymes (Kovalchuk et al. 2013; Kovalchuk and Driessen 2010; Víglaš and Olejníková 2021). Additionally, some white-rot fungi have the ability to use non-enzymatic processes to aid in wood degradation (Moretti et al. 2023)

Regardless, not all wood-rotting Basidiomycetes can be categorized as strictly ‘white-rot fungi’ or ‘brown-rot fungi’, based on the enzymes encoded by their genomes (or not) or results of various *in vitro* assays of wood degradation or enzyme activities (Riley et al. 2014). For example, the type of wood decay caused by *F. mediterranea* is characterized as a white rot, based originally on visibly descriptions of wood decay (Larignon and Dubos 1997), with further confirmation from *in vitro* assays of its enzymatic and non-enzymatic lignin-degrading properties (Cloete et al. 2015b; Moretti et al. 2023). In contrast, *F. polymorpha* has characteristics of both white-rot fungi (production of lignin-degrading enzymes) and brown-rot fungi (persistence of lignin in inoculated wood) (Galarneau et al., In review).

To advance our study of basidiomycetes that cause esca in grapevines, we have assembled high-quality genomes of the species *Fomitiporia mediterranea*, *Fomitiporia polymorpha*, *Tropicoporus texanus,* and *Inonotus vitis*. The type of wood-decay caused by the latter two species has not been characterized. Therefore, with the aim to describe the functional arsenal encoded by these set of species we performed functional annotation and comparative analysis, focusing on putative virulence factors in these newly sequenced genomes against those found in known species causing brown-rot, white-rot, in addition to ascomycete wood-decay fungi. Additionally, we identified a set of putative virulence factors undergoing expansion and contraction in these fungi, highlighting potential avenues for further research.

## Materials and Methods

### Sample collection

*Fomitiporia polymorpha*, *Inonotus vitis,* and *Tropicoporus texanus* were previously isolated in 2019 from grapevines (specifically *Vitis vinifera* L. ‘Chardonnay’ and *Vitis vinifera* ‘Cabernet-Sauvignon’) with the leaf symptoms of esca, in California, USA (Brown et al. 2020). *Fomitiporia mediterranea* was previously isolated in 1996 from grapevines (specifically *Vitis vinifera* ‘Ugni blanc’) with the leaf symptoms of esca, in the Charente wine region (France), and stored at the mycological collection at UMR SAVE (INRAE Nouvelle-Aquitaine Bordeaux) (Laveau et al. 2009).

### DNA extraction and sequencing

A pure isolate of *F. mediterranea* was grown in liquid malt medium (cristomalt 15 g/l) at 100 rpm for 22 days. Pure fungal isolates of *Fomitiporia polymorpha*, *Inonotus vitis,* and *Tropicoporus texanus* were grown in potato dextrose broth at 110 rpm for seven days. The mycelium was filtered and washed twice with sterile water using vacuum filtration. The dry mycelium was frozen in liquid nitrogen and ground with a TissueLyser II (Qiagen) using a 50-ml stainless-steel jar (Retch). High molecular DNA extraction was based on the method of Chin et al. (2016), using 5 g of ground mycelium. To confirm identity, we amplified and sequenced ITS (using primers ITS1-ITS4), elongation factor-1-alpha (using primers EF1-983F and EF1-1567R), and RPB2 (using primers bRPB2-6F and bRPB2-7.1R); the amplicons were aligned against NCBI nucleotide collection using blastn.

Prior to the preparation of the HiFi sequencing library of *F. mediterranea*, a preliminary step of short reads elimination (SRE, Pacific Biosciences, Menlo Park, CA, USA) was carried out. This process ensured the overall quality and integrity of the HMW gDNA before library prep. SRE pre-treated gDNA was sheared with a mode size of 15-18 Kbp using Diagenode’s Megaruptor (Diagenode LLC, Denville, NJ, USA). The HiFi SMRTbell template was prepared with 5 µg of sheared DNA using the SMRTbell Prep Kit 3.0 (Pacific Biosciences, Menlo Park, CA, USA) following the manufacturer’s instructions. Ampure PB bead size selection step was performed on HiFi SMRTbell template to deplete DNA fragments shorter than 5 Kbp using a 35 % v/v dilution of AMPure beads. Concentration and final size distribution of the library were evaluated using a Qubit 1X dsDNA HS Assay Kit (Thermo Fisher, Waltham, MA, USA) and Femto Pulse System (Agilent, Santa Clara, CA, USA), respectively. HiFi library was sequenced using a Revio sequencer (DNA Technology Core Facility, University of California, Davis).

The molecular weight of the DNA extracted from *Fomitiporia polymorpha*, *Inonotus vitis,* and *Tropicoporus texanus* was evaluated with pulse field electrophoresis (Pippin pulse, Sage Science). Two hundred microliters of HMW gDNA at a concentration of 100 ng/µl were fragmented using a 26G blunt needle (SAI Infusion Technologies). gDNA shearing was accomplished by aspirating the entire volume and passing the sample through the 26G blunt needle ten times. After shearing, the sample was cleaned and concentrated using 0.45X AMPure PB beads and the size distribution of the sheared gDNA fragments was assessed using pulse field gel electrophoresis (Pippin pulse, Sage Science, Beverly, MA, USA) prior to libraries preparation. SMRTbell templates wer prepared with 10 µg of sheared DNA using SMRTbell Express Template Prep Kit 2.0 (Pacific Biosciences, Menlo Park, CA, USA) following the manufacturer’s instructions. The Blue Pippin instrument (Sage Science, Beverly, MA, USA) was used to size select SMRTbell templates using a cut-off size of 20-80 Kb. Size-selected libraries were cleaned with 1x AMPure PB beads, and their concentration and final size distribution were evaluated using a Qubit HS Assay Kit (Thermo Scientific, Hanover Park, IL, USA) and pulse field gel electrophoresis (Pippin pulse, Sage Science) respectively. Finished libraries were sequenced on a Sequel II sequencer (DNA Technologies Core Facility, University of California, Davis).

### Genome assembly and polishing

The HiFi reads of *F. mediterranea* were assembled using Hifiasm v.0.19.5-r587 (Cheng et al. 2021). First, reads between 15 kbp to 25 kbp were selected. Multiple parameters with different values were tested to obtain the least fragmented genome. The optimized options were “-a 3 -k 41 -w 51 -f 0 -r 5 -s 0.8 -D 10 -N 10 -n 7 -m 10000000,” which generated the primary assembly and the haplotigs.

The subreads from the CLR sequencing of *F. polymorpha*, *I. vitis,* and *T. texanus* were randomly subsampled using bamsieve within the PacBio SMRT link v.8.0. using the option “--percentage 25” to keep only 25% of the reads. Canu v.2.2 (Koren et al. 2017) was used with the options “corOutCoverage=200 ‘batOptions=-dg 3 -db 3 -dr 1 -ca 50 -cp 50’ genomeSize=60m -pacbio” to assemble the genomes.

The pbmm2 v.0.8.1 align (https://github.com/PacificBiosciences/pbmm2) was used to map the subreads to the raw assembly using the options “--preset SUBREAD --sort -j 32 -J 8”. These alignments were used as input for gccp v2.02 (https://github.com/PacificBiosciences/gcpp) with the options “--algorithm=arrow -j 40” to polish the raw assembly. After confirming the duplication levels using BUSCO v.5.6.1 (Manni et al. 2021) with the basidiomycota_odb10 database, the genomes were split into primary scaffolds and the haplotigs. *Inonotus vitis* and *T. texanus* genomes were split using the pipeline purge_haplotigs (Roach et al. 2018) following the instruction at https://bitbucket.org/mroachawri/purge_haplotigs/src/master/. The purged genomes were evaluated with BUSCO to confirm their completeness and duplication levels.

The genome of *F. polymorpha* was tested further before continuing because purge_haplotigs yielded inconsistent results. First, the assembly was evaluated for external contamination using ContScout (Bálint et al. 2024) with the options “-d uniprotKB -a diamond -q 251363”. The Taxonomy id 251363 belonged to *F. polymorpha*. This analysis revealed no external contamination in the genome. Additionally, *in-silico* PCR of the ITS sequences in the genome was performed with in_silico_pcr.pl (https://github.com/egonozer/in_silico_pcr) with the primers ITS1 (5’-TCCGTAGGTGAACCTGCGG-3’) and ITS-4 (5’-TCCTCCGCTTATTGATATGC-3’). The amplicons were blasted against the NCBI nucleotide collection, and they mapped to *F. polymorpha*. Additional *in-silico* PCR was done with various bacterial markers; no amplificons were obtained. Next, the total predicted proteins of the species (See repeat and gene annotation) were used with OrthoFinder v.2.5.4 (Emms and Kelly 2019) with default parameters. This was done to obtain a gene copy number profile. The profile was compared to the profile of the diploid assembly of *F. mediterranea* and the results suggested an incomplete diploid assembly or a possible aneuploidy of *F. polymorpha*. Therefore, the primary assembly of the species was manually curated.

For the manual curation of *F. polymorpha*, the gene alignments were generated with blastall within BLAST+ 2.15.0 (Camacho et al. 2009) with the options “-p blastp -e 1e-10 -b 5 -v 5 -m 8 -a 8”. The alignments were used with the gff file of the genes to obtain the collinearity percentages using MCScanX (Wang et al. 2012) with default parameters. If more than 70% of the total genes in a scaffold were in collinearity with another scaffold, then it was moved to the haplotigs. This process allowed the curating of a primary assembly, retaining high completeness and low duplication based on the BUSCO analysis.

### Repeat and gene annotation

The primary assembly of the species was used to annotate and mask repeats. First, repeat model libraries were predicted for each genome using RepeatModeler v.1.0.8 (Smit et al. 2015b) with default parameters. RepeatMasker v.4.06 (Smit et al. 2015a) was used with default parameter and custom repeat libraries to annotate and mask the repeats. The custom library contained the predicted models and the repbase library 20160829-2023 (Bao et al. 2015). The repeat annotation was used to extract the Transposable Elements (TE) for further analysis. These TEs were classified into Class I and Class II.

The tool maskFastaFromBed (Quinlan 2014) was used with the option “-soft”, the repeat annotation, and the primary assembly to softmask the repeats. The genes were annotated using Braker v2.1.6 (Brůna et al. 2021). The soft-masked genomes with the OrthoDB11 database of fungi (Kuznetsov et al. 2023) were used with the options “--fungus --softmasking 1” to obtain the raw gene annotation. This annotation was further cleaned to remove proteins with internal stop codons and without stop or end codons.

To calculate the gene density, the tool makewindows from BEDtools v.2.29.1 (Quinlan 2014) was used to obtain windows of 10kb from the primary assemblies. Then, the coverage tool from the same package was used with the windows and the gene and repeat annotation to obtain the density of the feature per 10kb of genomic regions. These results were organized and plotted using ggplot2 v.3.5.0 (Wilkinson 2011) in R v.4.2.2 (R Core Team 2022). Similarly, in the TE composition plot, the elements that represented less than 1% of the total TE content in the species were grouped into the category others. The organized data was plotted with ggplot2.

### Functional annotation

The general annotation of the predicted proteins was assigned using Pfamscan (https://www.ebi.ac.uk/jdispatcher/pfa/pfamscan) against the Pfam-A database using an e value of 0.001. CAZymes were annotated with the dbCAN3 at https://bcb.unl.edu/dbCAN2/blast.php (Zheng et al. 2023), selecting the options “HMMER: dbCAN (E-Value < 1e-15, coverage > 0.35)”, “DIAMOND: CAZy (E-Value < 1e-102)” and “HMMER: dbCAN-sub (E-Value < 1e-15, coverage > 0.35)”. The annotation was kept only when the genes were annotated with at least two algorithms. The signal peptides were assigned using SignalP 5.0 (Almagro Armenteros et al. 2019). The proteins with annotation in both databases (SignalP5 and dbCAN3) were annotated as secreted CAZymes. Secondary metabolite clusters were annotated using antiSMASH v.6.0 at https://fungismash.secondarymetabolites.org with default parameters (Blin et al. 2021). The genes in each cluster were given annotations of the cluster to which they belonged. The peroxidases were annotated using a specialized database for fungi fPoxDB (Choi et al. 2014) using hmmsearch within HMMER v.3.1b2 (Eddy 2011) with the option “-E 1e-5”. The Cytochrome P450 proteins were annotated by intersecting the results of 3 methods, Pfam, blastp within Diamond v2.1.8.162 (Buchfink et al. 2021) with options “--id 60 --outfmt 6 --evalue 0.0001” against CYPED 6.0 database (Fischer et al. 2007), and phmmer (Eddy 2011) with options “-E 0.001 --incE 0.001 --noali” against CYPED 6.0 database. Genes with annotation with at least two methods were kept as P540. Last, the proteins involved in transportation functions were annotated using Diamond blastp with the option “-evalue 1e-10” against the TCDB database (Saier et al. 2021). The R package tidyheatmap v 0.2.1 (Mangiola and Papenfuss 2020) was used to build the heatmaps.

### Phylogenetic analysis

Genomes and gene annotations from the four Basidiomycete species were compared to those of eight Basidiomycete species and two Ascomycete species (**Supplementary Table 1**). As the four species are wood-colonizing fungi, we included for comparative purposes a combination of species that cause different types of wood decay, namely Basidiomycetes that cause brown rot or white rot and Ascomycetes that cause soft rot. White-rot fungi were *Pleurotus ostreatus*, *Stereum hirsutum,* and *Trametes versicolor* (**Supplementary Table 1**). Brown-rot fungi were *Daedalea quercina* (aka *Fomitopsis quercina*), *Fomitopsis schrenkii, Gloeophyllum trabeum*, *Postia placenta,* and *Serpula lacrymans*. Ascomycetes that causes the grapevine trunk disease Botryosphaeria dieback *Neofusicoccum parvum* (soft-rot fungus; Galarneau et al, In review) and *Botryosphaeria dothidea* were also included. Additionally, other non-pathogenic and pathogenic Ascomycetes known for colonizing wood and other organs of the plant were included for phylogenetic reference only (**Supplementary Table 1**).

The predicted proteins of all the genomes were used as input for OrthoFinder v.2.5.4 (Emms and Kelly 2019) with default parameters. The resulting single copy orthologs were aligned using MUSCLE v.5.1 (Edgar 2004) with the option “-maxiters 16”. The alignments were concatenated and parsed with Gblocks v.0.91b (Castresana 2000) with default parameters. ModelTest-NG v.0.1.7 (Darriba et al. 2020) was used to optimize the evolutionary model. The Maximum likelihood tree was obtained with RAxML-NG v.0.9.0 (Kozlov et al. 2019), the parsed alignment, and the evolutionary model “LG+I+G4” with the options “--tree pars{10} --bs-trees 100”. The clock-calibrated tree was constructed using BEAST v.2.7.6 (Bouckaert et al. 2019). The parsed alignment of single-copy orthologs was prepared with BEAUti v2.7.6 (Bouckaert et al. 2019). Calibration points were set for the Ascomycetes crown to 539 Mya (Prieto and Wedin 2013) with a normal distribution, and the Polyporales group set to 142 Mya (Ji et al. 2022) with a normal distribution. Five different Markov chain Monte Carlo chains of 1,000,000 generations were set. The LG substitution model with four gamma categories, a strict clock, and the Birth-Death model was used. Sampling was performed every 1,000 generations. The resulting log and tree files were combined using LogCombiner v.2.7.6 (Bouckaert et al. 2019), and the maximum clade credibility tree was generated using TreeAnnotator v2.7.6 (Bouckaert et al. 2019) with a burn-in of 10,000 generations. Figtree (Rambaut 2018) was used to plot the phylogenetic trees.

### Gene family expansion and contraction analysis

The predicted proteins in all the genomes were concatenated in a single file and blasted to themselves using blastp within Diamond v2.1.8.162 (Buchfink et al. 2021) with the options “--evalue 1e-6 --very-sensitive --outfmt 6”. The proteins were grouped in families using Markov clustering with MCL v.14-137 (Enright et al. 2002). To do the clustering, first mcxload with the options “--stream-mirror --stream-neg-log10 -stream-tf ‘ceil(200)’” was used with the blast result file. Next, the output from mcxload was used as input for mcl with the option “-I 3”. Then, mcxdump was used with the output from the previous step to obtain a preliminary file needed to run CAFE v.5.0 (Mendes et al. 2021). The script cafetutorial_mcl2rawcafe.py at https://github.com/hahnlab/cafe_tutorial/tree/main/python_scripts was used to prepare the file. CAFE was run with the option “-P 0.0100”, an estimated lambda value of 0.0011169463915058, the previously prepared input, and the clock-calibrated tree. The resulting file was used to extract the families with significant rates of gain or loss of genes (*P* value < 0.01). The number of families expanding and contracting per phylogenetic node was incorporated into the clock-calibrated tree using CafePlotter (https://github.com/moshi4/CafePlotter). A Fisher’s exact test was used to obtain the functions that were significantly enriched in the expanded and contracted families of the species of interest.

### Data availability

The sequencing data generated for this project are available at NCBI (BioProject: PRJNA1099290). The references for previously published genomic data used in this study can be seen in **Supplementary Table 1**. The genome assemblies and gene models produced in this study are publicly available at Zenodo (https://doi.org/10.5281/zenodo.10957629). Dedicated genome browsers and BLAST tools are available at https://www.grapegenomics.com. The code used in this project is displayed in **Supplementary File 1**.

## Results

### Genome assemblies of Fomitiporia mediterranea, Fomitiporia polymorpha, Inonotus vitis, and Tropicoporus texanus

The isolates of the four Basidiomycete species described in this study were obtained from esca-symptomatic plants in vineyards in US states California and Texas (Brown et al. 2020), and in Charente in France (Laveau et al. 2009). The total genome assembly sizes were 148.9 Mbp for *F. polymorpha*, 126.2 Mbp for *F. mediterranea*, 71.7 Mbp for *I. vitis*, and 68.3 Mbp for *T. texanus*. The genome coverage ranged from 40 X in *F. polymorpha* to 148 X in *F. mediterranea*. Except for *F. polymorpha*, all genomes were divided into a primary assembly and haplotigs, each constituting roughly half the size of the total genome assembly (**Table 1**). This even distribution between the primary assembly and haplotigs suggests the complete diploid representation of the genomets. Conversely, the *F. polymorpha* assembly underwent manual curation, resulting in a primary assembly of 85.6 Mbp and a set of haplotigs totaling 63.2 Mbp. After ruling out the possibility of contamination with DNA from another species in the sequencing reads, a gene copy number analysis for *F. polymorpha* and *F. mediterranea* revealed a higher proportion of odd copy numbers in *F. polymorpha* compared to the diploid genome of *F. mediterranea* (**Supplementary Figure 1**). This finding suggests that either the diploid assembly of the *F. polymorpha* genome was incomplete or aneuploid.

**Table 1.**
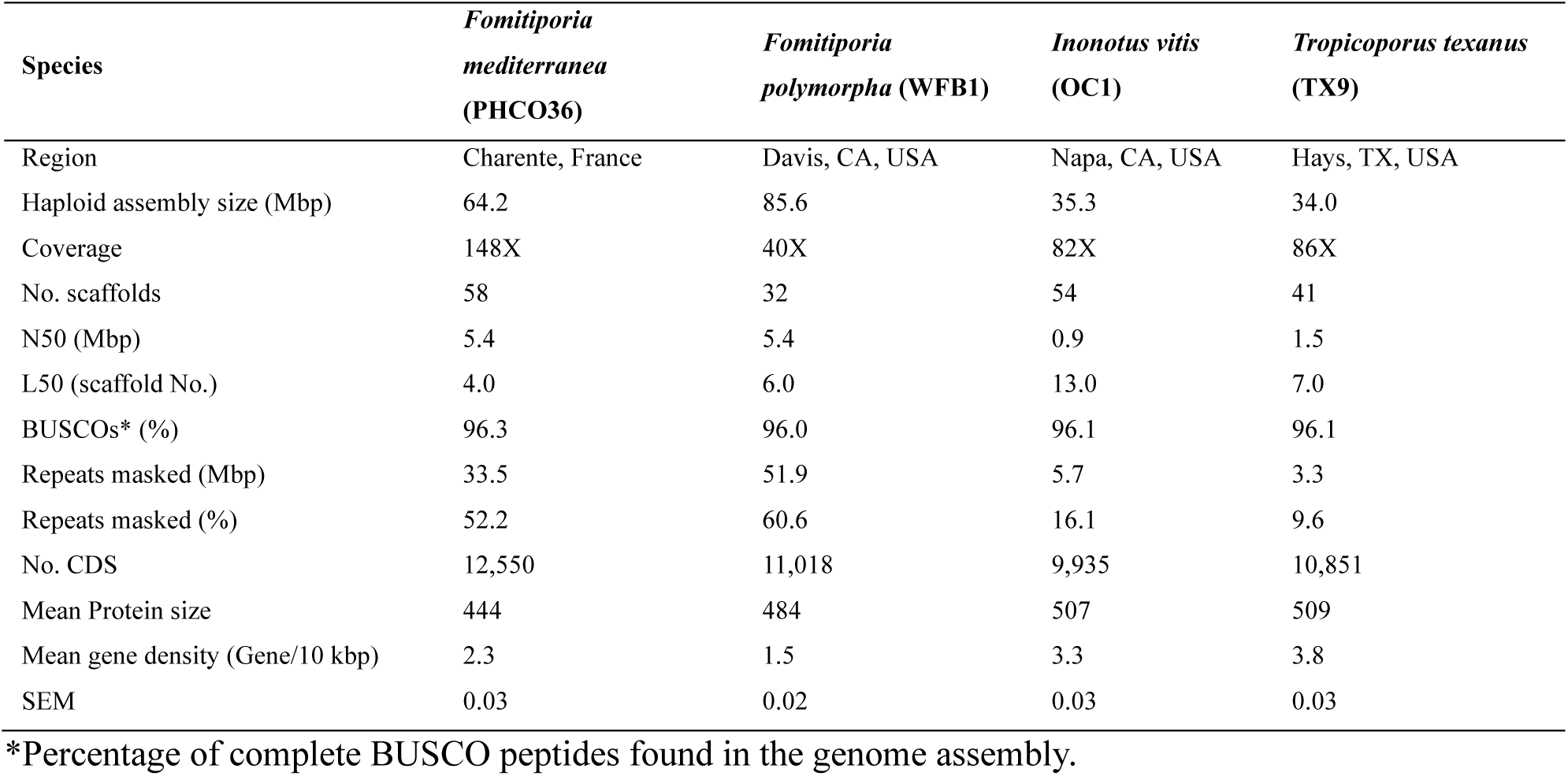
Assembly and annotation statistics of the four basidiomycete species genomes.

The statistics presented henceforth pertain to the primary assembly (i.e., the more contiguous haploid versions) of the genomes. The number of scaffolds in the haploid genomes varied from 32 in *F. polymorpha* to 58 in *F. mediterranea* (**Table 1**). The Benchmark of Universal Single-Copy Orthologues (BUSCO) analysis showed that the assemblies contain over 96% of the complete single-copy ortholog sequences curated for the basidiomycetes group (**Table 1**). The highest repeat content was observed in *F. polymorpha*, constituting more than 60% of the genome size, followed by 52.2% in *F. mediterranea*, 16.1% in *I. vitis*, and 9.6% in *T. texanus*. This distribution of repeat content aligns with gene density, with the highest density observed in *T. texanus* and the lowest in *F. polymorpha* (**Table 1**). Comparing the density of genes and repeats within species revealed that repeats were significantly denser in *F. polymorpha* and *F. mediterranea*. Conversely, in *I. vitis* and *T. texanus*, the genes were significantly denser than the repeats (**Figure 1A**). Additionally, a comparison of feature density between species showed that *I. vitis* and *T. texanus* had significantly higher gene density compared to both *Fomitiporia*. The reverse was true for repeat content, where both *Fomitiporia* species had higher repeat-content density than *I. vitis* and *T. texanus* (**Figure 1B**).

**Figure 1.**
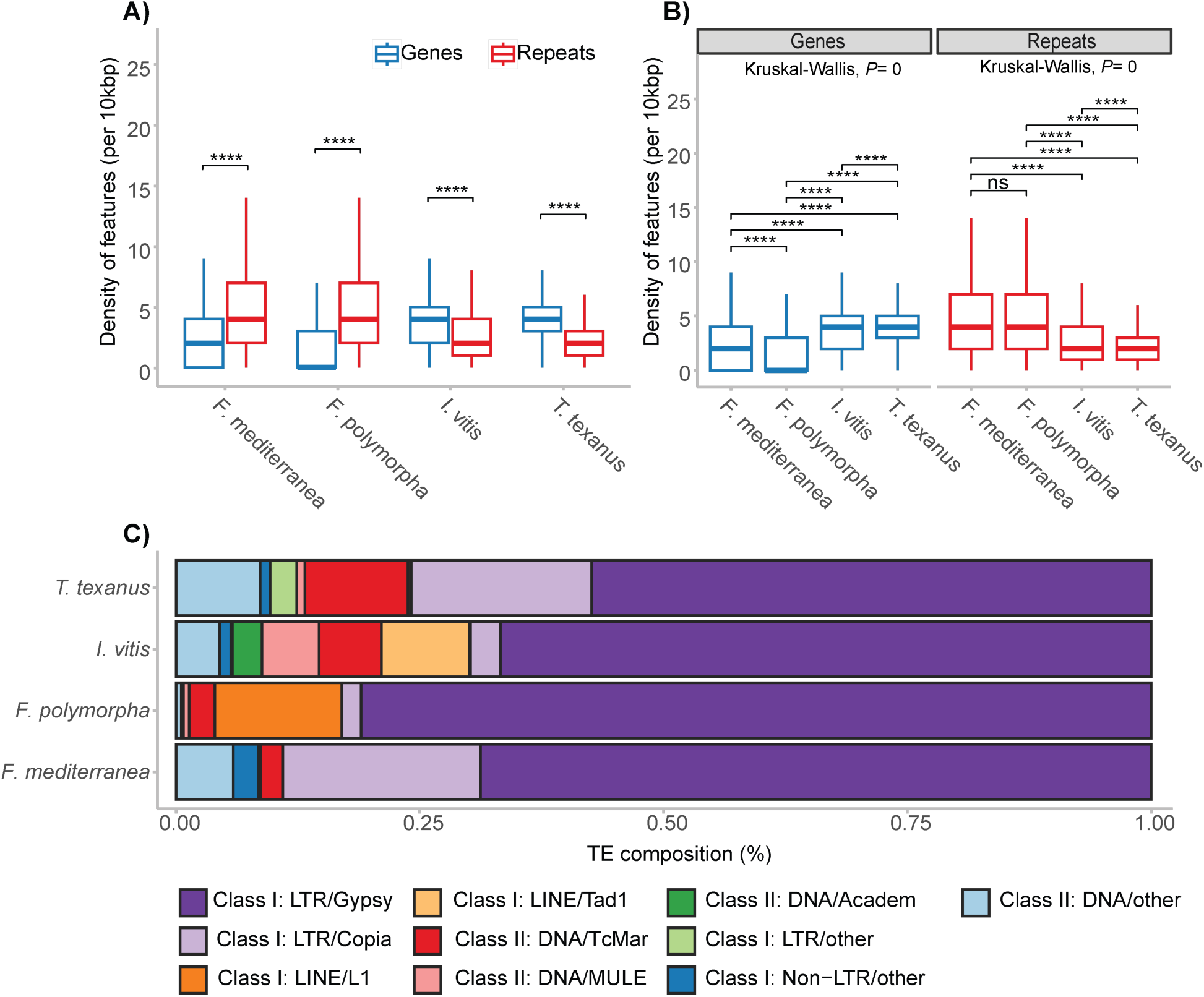
Genes and repeats density in the genome of the species in study. **A**, Density of genes and repeats per 10 kbp of genome space with statistical comparisons within species. Significant differences were determined using the Mann-Whitney U test (**** *P* ≤ 0.0001). **B**, Density of genes and repeats per 10 kbp of genome space with statistical comparisons between species. Significant differences were determined using the Dunn’s test after a significant Kruskal-Wallis test (**** *P* ≤ 0.0001). **C,** Transposable element (TE) types as a proportion of the total TE length in the genome. Categories representing < 1% of the repeat content were classed as “other.”

A significant proportion of total repeats were transposable elements (TEs), with 48% of the total repeat content in *T. texanus*, 68% in *F. mediterranea*, 71% in *I. vitis*, and 75% in *F. polymorpha*. Within the TEs, the LTR/Gypsy covered an average of 68.5 ± 4.9%, the most significant proportion of the TEs. *F. polymorpha* presents the largest proportion, with 81.0% of their total TEs sequence being LTR/Gypsy (**Figure 1C**). Next, the LTR/Copia elements covered a large proportion of TEs in *F. mediterranea* (20.3%) and *T. texanus* (18.5%). Moreover, the elements LINE/Tad1 and DNA/MULE were almost exclusive to *I. vitis*, whereas the LINE/L1 was almost exclusive to *F. polymorpha* (**Figure 1C**).

### Gene annotation focused on putative virulence factors

We focused the functional annotation of the protein-coding genes of three species on the potential virulence factors such as Carbohydrate-Active Enzymes (CAZymes), fungal peroxidases, cytochrome P450s, biosynthetic gene clusters (BGCs), and cellular transporters. The variation in gene numbers among putative virulence factors can indicate the virulence strategies adopted by fungi. For example, in previous studies among the species that cause Botryosphaeria dieback, the more virulent *Neofusicoccum* species have a larger repertoire of carbohydrate-active enzymes (CAZymes), compared to those of less virulent *Dothiorella* spp. and *Diplodia* spp. (Garcia et al. 2021). Similarly, our observations reveal that grapevine trunk pathogens like *Eutypa lata* and *Phaeoacremonium minimum* exhibit a higher number of biosynthetic gene clusters (BGCs) than *N. parvum* (Garcia et al. 2024). These observations are supported by evidence of recent evolutionary changes like gene family expansions in these virulence factor groups (Garcia et al. 2021, 2024).

In this study between the Basidiomycete species, both *Fomitiporia* genomes encoded a high number of fungal peroxidases, similar to those of other white-rot species, such as *Trametes versicolor. T. texanus* encoded a high number of secreted CAZymes, whereas *I. vitis* appeared to be on the medium or lower end of counts per functional category when compared with all the species (**Table 2**).

**Table 2.** Genes annotated per functional category.

| Species | Total predicted genes | Signal peptide | CAZymes | Secreted CAZymes | Peroxidases | P450 | BGC genes | Transporters |
| --- | --- | --- | --- | --- | --- | --- | --- | --- |
| *Saccharomyces cerevisiae* | 6,716 | 352 | 137 | 41 | 21 | 3 | 31 | 1,670 |
| *Botryosphaeria dothidea* | 12,424 | 1,341 | 611 | 348 | 56 | 262 | 926 | 3,219 |
| *Neofusicoccum parvum* | 13,067 | 1,429 | 642 | 390 | 58 | 272 | 930 | 3,363 |
| *Fomitiporia mediterranea* | 12,550 | 888 | 408 | 233 | 51 | 137 | 330 | 2,597 |
| *Fomitiporia polymorpha* | 11,018 | 911 | 429 | 247 | 55 | 154 | 305 | 2,736 |
| *Inonotus vitis* | 9,935 | 801 | 377 | 220 | 47 | 118 | 299 | 2,357 |
| *Tropicoporus texanus* | 10,851 | 925 | 440 | 268 | 49 | 114 | 336 | 2,488 |
| *Stereum hirsutum* | 14,072 | 1,059 | 521 | 294 | 53 | 211 | 408 | 2,711 |
| *Gloeophyllum trabeum* | 11,846 | 779 | 345 | 180 | 32 | 123 | 480 | 2,417 |
| *Pleurotus ostreatus* | 12,330 | 1,094 | 507 | 283 | 54 | 151 | 295 | 2,450 |
| *Serpula lacrymans* | 12,789 | 686 | 297 | 144 | 35 | 159 | 609 | 2,339 |
| *Trametes versicolor* | 14,296 | 1,084 | 450 | 257 | 68 | 188 | 428 | 2,499 |
| *Postia placenta* | 12,541 | 737 | 276 | 114 | 37 | 215 | 597 | 2,359 |
| *Daedalea quercina* | 12,199 | 790 | 301 | 152 | 34 | 138 | 389 | 2,386 |
| *Fomitopsis schrenkii* | 13,885 | 893 | 351 | 177 | 37 | 182 | 297 | 2,509 |

The abundance patterns of specific families within the functional categories varied between the species analyzed. Among the families of the Cytochrome P450 enzymes, the activity can range from synthesizing secondary metabolites to degrading xenobiotic molecules, which generally contribute to adaptation to different environmental conditions (Chen et al. 2014; Moktali et al. 2012; Črešnar and Petrič 2011). Interestingly, the families CYP620 and CYP512 appeared to be exclusive to basidiomycete species. These families are involved in the polyketide biosynthesis (Yu et al. 2020; Nazmul Hussain Nazir et al. 2010) and triterpenoid biosynthesis (Yu et al. 2020; Syed et al. 2014; Chen et al. 2012), respectively. The number of genes annotated as CYP512 was particularly high in *F. polymorpha*. Similarly, CYP53, involved in detoxifying benzoate residues (Faber et al. 2001), was highly abundant in *F. polymorpha* (**Figure 2**).

**Figure 2.**
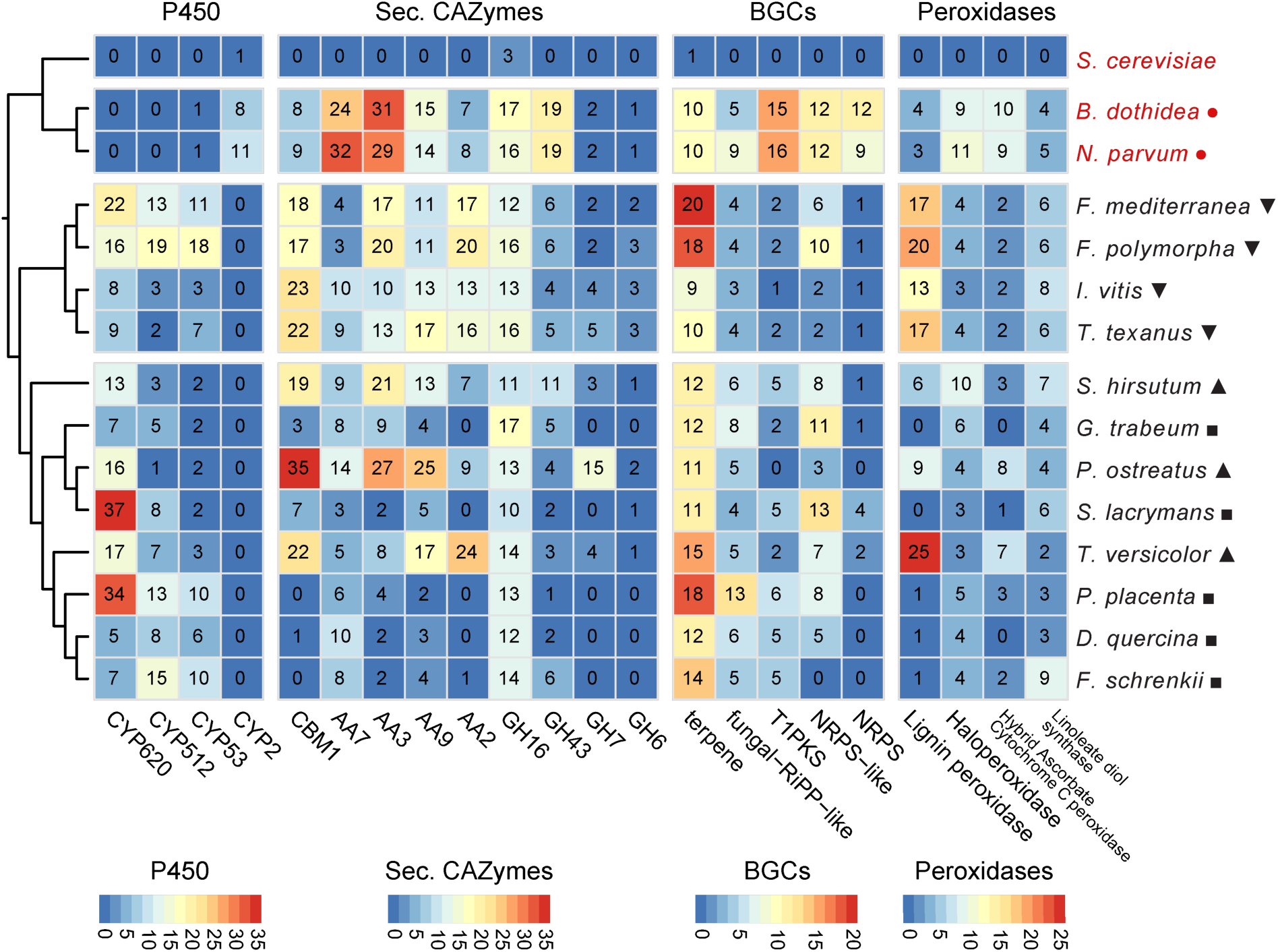
Number of protein-coding genes annotated as putative P450, secreted CAZymes, BGC, and peroxidases. The specific functions per functional category were selected based on the highest number of genes across all genomes. Ascomycete species are written in red, and Basidiomycete species in black. Species .marked with ▾ are the fungi of interest in this study, ▴ are known white-rot species, ▪ are known brown-rot species and ● are known soft rot species.

The successful establishment of these fungi in their hosts relies heavily on their ability to degrade the various polysaccharides that constitute the plant cell wall (Morales-Cruz et al. 2015). CAZymes are primarily responsible for the modification of these host cell wall polysaccharides (Kubicek et al. 2014). The identification of CAZymes with signal peptides for secretion is a common method for studying the potential array of enzymes capable of degrading the plant cell wall (Garcia et al. 2021; Blanco-Ulate et al. 2014; Morales-Cruz et al. 2015). Compared to white-rotters *S. hirsutum* and *T. versicolor,* the species *F. mediterranea*, *F. polymorpha, I. vitis*, and *T. texanus* were characterized by similarly high number of genes encoding Carbohydrate-Binding module 1 (CBM1), which is associated with cellulose-binding activity (Lehtiö et al. 2003). In contrast, CMB1 was absent from brown-rotters *P. placenta* and *F. schrenkii* (**Figure 2**). The Auxiliary Activity Families (AA) 7, 3, and 9 are gluco-oligosaccharide oxidases, cellobiose dehydrogenase, and polysaccharide monooxygenases, respectively. AA7 and AA3 were more prevalent in the pathogens *B. dothidea* and *N. parvum*, whereas AA3 also showed high gene counts in *P. ostreatus*, *S. hirsutum*, *F. polymorpha*, and *F. mediterranea*. Conversely, AA9 was more abundant in species like *P. ostreatus*, *T. texanus*, and *T. versicolor* (**Figure 2**). The members of the Glycoside hydrolase family 16 (GH16) are widely distributed across different taxonomic groups. Their primary functions include the degradation or remodeling of cell wall polysaccharides, such as those involving endo-glucanases, transglycosylases, and chitin-glucosyltransferase (Viborg et al. 2019). This gene family is consistently large across all the species studied, except for *S. cerevisiae*, where only three genes have been annotated as GH16 (**Figure 2**). Additionally, members of GH43, known for their xylosidase and arabinofuranosidase activities (Flipphi et al. 1993; Shallom et al. 2005), were found to be more abundant in *B. dothidea* and *N. parvum*. The glycoside hydrolase families GH7 and GH6, with cellobiose hydrolase functions (Riley et al. 2014), were almost absent from the brown-rot species. On the other hand the *F. mediterranea*, *F. polymorpha*, *I. vitis* and *T. texanus* have similar number of GH7 and GH6 than those of the white-rot species.

Biosynthetic gene clusters (BGCs) produce secondary metabolites critical for fungal development and interactions with their plant host and other organisms (Keller 2019). *F. mediterranea*, *F. polymorpha*, and *P. placenta* showed the highest number of terpene BGCs. Compared to all brown-rotters and all white-rotters, *F. mediterranea*, *F. polymorpha I. vitis*, and *T. texanus* were characterized by similarly low numbers of type I Polyketide synthases (T1PKS) and Non-Ribosomal Peptide Synthetases (NRPS). In contrast, T1PKS and NRPS were most numerous in soft-rotter *N. parvum* and related species *B. dothidea*. Additionally, as previously mentioned, the capacity to fully degrade lignin significantly distinguishes white-rot from brown-rot basidiomycetes. White-rot species *T. versicolor*, *P. ostreatus*, and *S. hirsutum*, along with *F. mediterranea*, possessed a significantly higher number of lignin peroxidases compared to brown rot species (*P=0.049*; **Figure 2**). However, the new genomes of *F. polymorpha*, *I. vitis,* and *T. texanus* showed no significant difference with the aforementioned white-rot fungi (*P*= 0.015).

### Gene family expansion and contraction

Next, we tested if the differences in putative virulence factor repertoires reflected accelerated processes of gene family expansion and contraction. We implemented the Computational Analysis of gene Family Evolution (CAFE) approach (Mendes et al. 2021). CAFE computes the birth and death rates of genes within gene families, identifies families with accelerated rates of gain and loss, and pinpoints the specific lineages responsible for significant changes in family size. CAFE requires a clock-calibrated phylogenetic tree and the sizes of gene families as input.

To establish the phylogenetic relationships of the species under study, we obtained a set of single-copy orthologs for all species (**Supplementary Table 1**) using Orthofinder. These orthologs were used to build a Maximum Likelihood (ML) tree (**Supplementary Figure 2**), with most tree nodes showing a bootstrap value of 100, indicating high topological accuracy. The newly presented genomes in this study form a single clade, clustering the *Fomitiporia* species and *I. vitis* with *T. texanus*. Upon confirming the ML tree’s topology, we used the set of orthologous genes to generate a clock-calibrated tree through a Bayesian approach with BEAST (Bouckaert et al. 2019). The crown ages of the Ascomycota and the Polyporales groups served as calibration points (**Figure 3A**). The topology of the Bayesian tree aligns with that of the ML tree, and the clock calibration suggests that the four studied species shared a common ancestor approximately 80 million years ago (Mya). The *Fomitiporia* species diverged around 8 Mya, while *I. vitis* and *T. texanus* diverged approximately 55 Mya.

**Figure 3.**
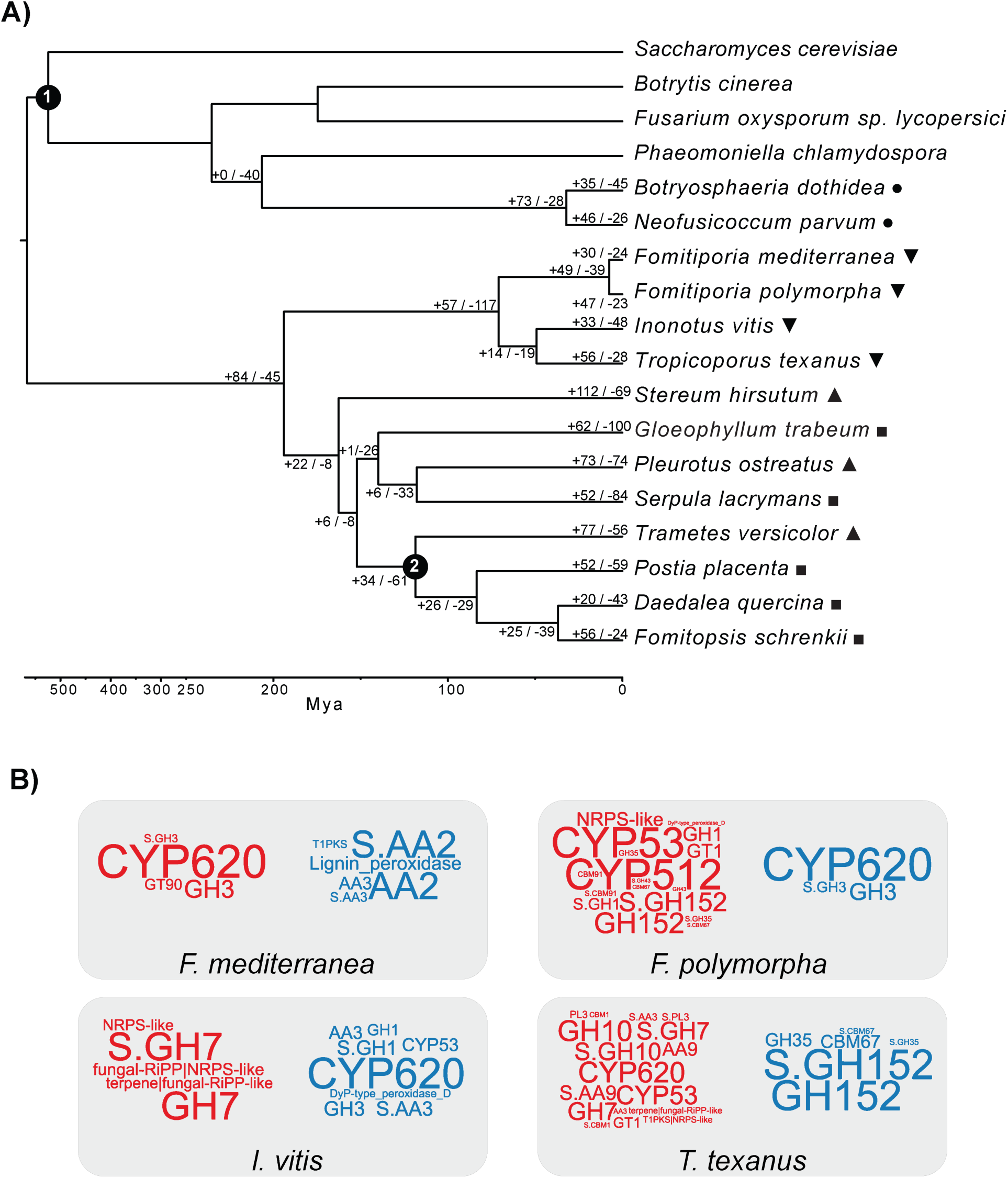
Analysis of gene family expansion and contraction in the species of interest. **A,** Clock calibrated phylogenetic tree constructed with single-copy gene orthologues. The branches represent divergence times in million years. Calibration point **(1)** at Ascomycetes crown set to ∼539 Mya. Calibration point **(2)** at the Polyporales group set to ∼142 Mya. Positive and negative numbers indicated on the branches represent expansions and contractions, respectively, as determined by gene family evolution analysis CAFE. Species marked with ▼are the fungi of interest in this study, ▲ are known white-rot species, ▪ are known brown-rot species, and ● are soft-rot Ascomycete species. The unmarked species were included for phylogenetic reference only. **B**, word clouds representing the genes enriched in the rapidly evolving families of the species *Fomitiporia mediterranea, Fomitiporia polymorpha*, *Inonotus vitis*, and *Tropicoporus texanus*. The word size represents the enrichment’s strength based on the *P* value. The red color represents the enriched genes within expanding families, and the blue represents enriched genes within contracting families.

The clock-calibrated tree, along with the gene family sizes for all species, served as input for CAFE. *F. polymorpha*, *F. mediterranea,* and *T. texanus* showed an overall expansion of gene families. Conversely, in *I. vitis,* the majority of the gene families are contracting (**Figure 3A**). An enrichment test conducted on the genes from families with accelerated gain or loss rates of gain or loss identified specific functions associated with these rapidly evolving families (**Figure 3B**).

Genes annotated as CYP620 were highly enriched in the expanded gene families of *F. mediterranea* (**Figure 3B**), whereas lignin peroxidases and AA2 were enriched in the contracting families. In *F. polymorpha*, CYP512, CYP53, and secreted GH152 showed high enrichment in expanding families (**Figure 3B**). Similarly, in *I. vitis*, genes of the BGC fungal-RiPP|NPRS-like and the secreted GH7 were enriched in expanding families (**Figure 3B**), whereas CYP620 was highly enriched in the contracting families. Lastly, in *T. texanus*, genes of the secreted CAZymes (GH10 and GH7) and P450s (CYP620 and CYP53) were enriched in the expanding families, while GH152 genes were highly enriched in the contracting families (**Figure 3B**).

## Discussion

Only a few Basidiomycete species have been associated with grapevine diseases and predominantly with esca. Traditionally, these species have been considered secondary pathogens, as they seemingly lack the capability to independently cause disease symptoms. However, researchers have observed a reduction of esca leaf symptoms after removing the white rot through vine surgery (Lecomte et al. 2022). More recently, researchers in California have demonstrated that some species can, in fact, trigger and exacerbate esca-related symptoms (Brown et al. 2020). In our study, we have assembled the genomes of Hymenochaetaceae family members associated with esca across various viticulture regions worldwide. With the exception of *Fomitiporia mediterranea* (Floudas et al. 2012), these genome assemblies are the first to be published for each species.

The haploid genome sizes of previously published genomes in the Hymenochaetaceae family range from approximately 28 Mbp to 63 Mbp. This range encompasses the assembly sizes of *I. vitis* and *T. texanus* presented in this study. The manually purged assembly size of *F. polymorpha* exceeds that of its close relative, *F. mediterranea*, by roughly 20 Mbp. Nonetheless, the completeness and duplication levels, assessed with BUSCO, are comparable to those of the other assemblies. Identifying more than 96% of the Basidiomycete orthologs in our assemblies aligns with studies on close relatives, where completeness varied from 84.1% to 94.9% (Zhao et al. 2023a). Additionally, the low number of scaffolds (from 32 to 58), the low L50 values (from 4 to 13), and the high N50 values (from 0.9 Mbp to 5.4 Mbp) of our assemblies indicate highly contiguous genomes, suggesting the presence of complete and nearly complete chromosomes. It is worth nothing that the sequencing technology and assembly software likely had an effect on the coverage differences observed between the species, where *Fomitiporia mediterranea*, the only species sequenced with HiFi presented higher coverage.

The repetitive content in fungal genomes directly affects genome plasticity, including both structural and functional alterations (Castanera et al. 2017). Repeat coverage values exceeding 41% have been documented for *F. mediterranea* (Floudas et al. 2012) and 21% *I. obliquus* (Duan et al. 2022), aligning with the findings for our genomes. As expected, the high repetitive content observed in Fomitiporia species corresponds to a sparser gene space compared to other genomes analyzed in this study. Among the identified repetitive sequences in the species of interest, LTR/Gypsy and LTR/copia elements were the most prevalent, a finding that is consistent with other fungal studies (Floudas et al. 2012; Castanera et al. 2017, 2016). Moreover, the presence and coverage of elements such as LINE/L1 and LINE/Tad1 in *F. polymorpha* and *I. vitis* are of particular interest. Despite various studies indicating that these elements are not uncommon in fungi, they are typically present in small quantities (Castanera et al. 2017, 2016; Qian et al. 2023; Daboussi and Capy 2003).

Annotation of the genes in the newly assembled genomes is comparable with those of other fungal pathogens, as reported in **Table 2**, ranging from ∼10k to ∼12k genes. Besides the widely known Ascomycetes pathogenic species *N. parvum* and *B. dothidea*, the new genomes of *F. mediterranea*, *F. polymorpha,* and *T. texanus* present a high number of genes annotated as putative virulence factors, which support the idea of their role as important wood decay microorganisms. More specifically, the P450 gene family CYP53 is highly abundant in *F. polymorpha*. This family confers abilities related to the degradation of benzoate residues, which is a common plant defense compound (Podobnik et al. 2008; Lah et al. 2011; Jawallapersand et al. 2014). Similarly, the CBM1, AA9, and AA2 CAZymes, higher in white-rot fungi and the newly assembled genomes, are involved in lignocellulose substrate degradation (Lehtiö et al. 2003; Liu et al. 2017; Garrido et al. 2020), suggesting these enzymes may play a crucial role in the activity of the white-rot fungi, consistent with previous reports by other researchers (Liu et al. 2017; Wu et al. 2020; Li et al. 2019; Miyauchi et al. 2020; Levasseur et al. 2014; Floudas et al. 2020). Additionally, researchers have reported high correlation between white-rot species and the families CBM1, GH7 and GH6 (Riley et al. 2014; Floudas et al. 2015, 2012). This groups of CAZymes were also observed in the newly assembled genomes in this study. The same applies to lignin peroxidases observed in the white-rot pathogens and the newly assembled genomes. Moreover, besides species like *B. dothidea* and *N. parvum* encoding more polyketide synthase clusters, more terpene clusters were annotated in the *Fomitiporia* species. The production of terpene metabolites like frustulosin and dihydroactinolide have been reported in *F. mediterranea* (Fischer and Thines 2017), and multiple terpene synthases have been annotated in the close relative *Inonotus obliquus* (Duan et al. 2022).

We assessed the relationship between the species presented in this study through maximum likelihood and Bayesian phylogenetic trees of single-copy orthologs. The separation of the ascomycetes from the basidiomycetes, the clustering of the members of the Hymenochaetaceae family (*F. mediterranea*, *F. polymorpha*, *I. vitis,* and *T. texanus*), and the grouping of the members of the order Polyporales are consistent with the literature (Ji et al. 2022; Song and Cui 2017; Zhao et al. 2023b; Chen et al. 2015; Liu et al. 2023). Additionally, the clock-calibrated tree estimation for the split between the *Fomitiporia* species is consistent with other studies (Chen et al. 2015; Liu et al. 2023).

The clock-calibrated tree allowed us to study patterns of gene family expansion and contraction in the newly assembled genomes. Identifying gene families with higher rates of gain and loss can help elucidate how these fungi may be adapting to their environments (Hahn et al. 2005). The enrichment of CYP620 in the expanding families of *F. mediterranea* suggests that the production of polyketides, a class of secondary metabolites with a wide range of activities, may play an important role in this species (Macheleidt et al. 2016). Also, the lignin peroxidases are enriched in the contracting families of *F. mediterranea*, an unexpected result based on their known ability to degrade lignin (Schilling et al. 2022). This result indicates that the common ancestor probably encoded a larger set of lignin peroxidases, an idea supported by the fact that the species *F. polymorpha* (phylogenetically closer to the common ancestor (Alves-Silva et al. 2020) encodes more proteins with this function than *F. mediterranea*.

Multiple gene families in *F. polymorpha* are undergoing expansion, and genes like CYP512, CYP53, and secreted GH152 are enriched in these families. As mentioned earlier, these P450 genes are involved in producing secondary metabolites needed for interaction with the environment and the degradation of benzoate produced by plants, improving fungal defense mechanisms. On the other hand, GH152 presents endo-β-1,3-glucanase activity, which targets fungal cell walls (Matsumoto et al. 2023; Gavande and Goyal 2023; Grenier et al. 2000). The role of this enzyme in fungi is not fully understood, but studies have suggested activity in modifying fungal cell walls and senescence of fruiting bodies (Grenier et al. 2000; Sakamoto et al. 2006), which could be related to competition with other fungi.

*Inonotus vitis* and *T. texanus* showed a similar pattern of expansion and contraction when compared to the *Fomitiporia* species. The significant enrichment in secondary metabolite biosynthetic genes, glycoside hydrolases, and P450s in the expanding families suggests that the repertoire of virulence factors of these pathogens has recently evolved, possibly in response to pressure imposed by the plant hosts, the environment, or microbial competitors and antagonists (Sacristán et al. 2021; Gladieux et al. 2014). Intraspecific genome comparisons would help determine whether these gene families are still evolving. The expansion of secreted GH7 and GH10 suggests a potential increase in cellulose and hemicellulose degradation capabilities. Similarly, the expansion of CYP53 implies evolution driven by the pressure imposed by plant defense compounds. In summary, these four basidiomycetes are actively evolving various functions of their putative virulence arsenal, some of which could improve their ability to thrive in their role as wood decomposers.

## Acknowledgments

This work was supported by the USDA National Institute of Food and Agriculture Specialty Crop Research Initiative (grant 2012-51181-19954). D. Cantu was also partially supported by the Louis P. Martini Endowment in Viticulture. D. Cantu, K. Baumgartner, and C.E.L Delmas by the Champion program framework INRAE – UC Davis. We thank the DNA Technologies Core at the Genome Center (UC Davis), and Dr. Andrea Minio (UC Davis), Nathalie Ferrer and Olivier Fabreguettes (INRAE) for their technical support.

## Supplementary files

**Supplementary Table 1.** Isolate and reference of the species used for the phylogeny and comparative analysis.

| Species | Isolate | Reference |
| --- | --- | --- |
| *Saccharomyces cerevisiae* | S288C-R64-4-1_20230830 | https://www.yeastgenome.org |
| *Botrytis cinerea* | v1.0 | (Amselem et al. 2011) |
| *Fusarium oxysporum* f. sp. *lycopersici* | 4287 v2 | (Ma et al. 2010) |
| *Phaeomoniella chlamydospora* | UCRPC4 | (Morales-Cruz et al. 2015) |
| *Botryosphaeria dothidea* | 0053 | (Garcia et al. 2021) |
| *Neofusicoccum parvum* | UCD646So | (Massonnet et al. 2018) |
| *Fomitiporia mediterranea* | PHCO36 | (Laveau et al. 2009), this study |
| *Fomitiporia polymorpha* | WFB1 | (Brown et al. 2020), this study |
| *Inonotus vitis* | OC1 | (Brown et al. 2020), this study |
| *Tropicoporus texanus* | TX9 | (Brown et al. 2020), this study |
| *Stereum hirsutum* | FP-91666 SS1 v1.0 | (Floudas et al. 2012) |
| *Gloeophyllum trabeum* | v1.0 | (Floudas et al. 2012) |
| *Pleurotus ostreatus* | PC15 v2.0 | (Riley et al. 2014) |
| *Serpula lacrymans* | S7.9 v2.0 | (Eastwood et al. 2011) |
| *Trametes versicolor* | v1.0 | (Floudas et al. 2012) |
| *Postia placenta* | MAD-698-R-SB12 v1.0 | (Gaskell et al. 2017) |
| *Daedalea quercina* | v1.0 | (Nagy et al. 2016) |
| *Fomitopsis schrenkii* | FP-58527 SS1 v3.0 | (Floudas et al. 2012) |

**Supplementary Figure 1.**
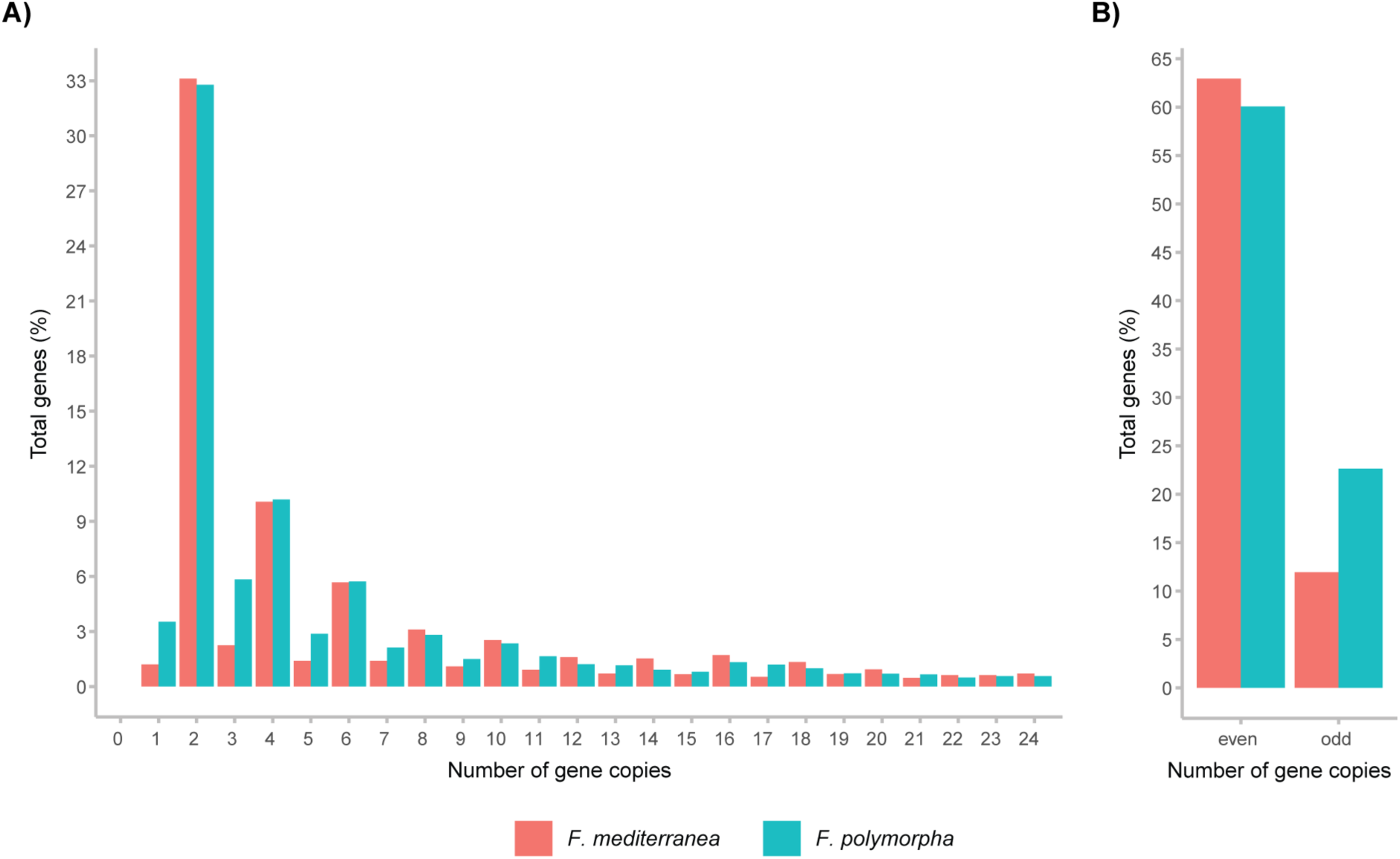
Gene copy number analysis of *F. mediterranea* and *F. polymorpha*. The figure shows a comparison of the gene copy number in the diploid genome of *F. mediterranea* and the full assembly of *F. polymorpha*. **A),** proportion of total genes per species with different copy numbers. **B)**, proportion of total genes per species with even or odd copy numbers.

**Supplementary Figure 2.**
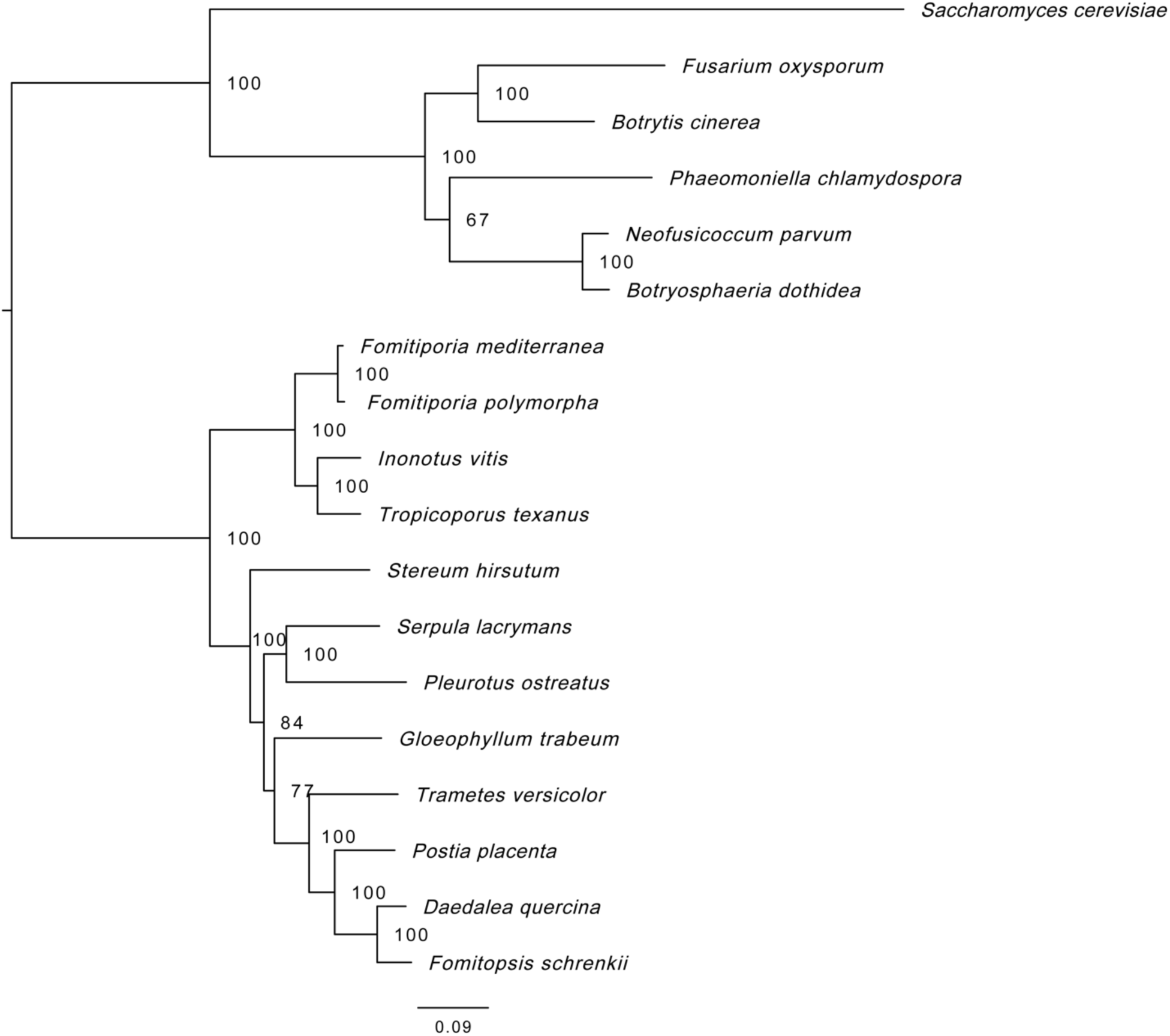
Maximum likelihood phylogenetic tree of the species in the study. The tree was constructed with a set of single-copy orthologs of the species in the study.

## Notes

https://www.grapegenomics.com

### Competing Interest Statement

The authors have declared no competing interest.

